# *Arabidopsis* MADS-box transcription factor AGL21 acts as environmental surveillance for seed germination by regulating *ABI5*

**DOI:** 10.1101/082164

**Authors:** Lin-Hui Yu, Jie Wu, Zi-Qing Miao, Ping-Xia Zhao, Zhen Wang, Cheng-Bin Xiang

## Abstract

Seed germination is a crucial checkpoint for plant survival under unfavorable environmental conditions. Abscisic acid (ABA) and its signaling play a vital role in integrating environmental information to regulate seed germination. MCM1/AGAMOUS/DEFICIENS/SRF (MADS)-box transcription factors are mainly known as key regulators of seed and flower development in *Arabidopsis*. However, their functions in seed germination are still poorly understood. Here we report that MADS-box transcription factor AGL21 negatively modulates seed germination and post-germination growth by controlling the expression of *ABA-INSENSITIVE 5* (*ABI5*) in *Arabidopsis. AGL21* responds to multiple environmental stresses and plant hormones. The *AGL21*-overexpressing plants are hypersensitive to ABA, salt and osmotic stresses during seed germination and early post-germination growth, whereas *agl21* mutants are less sensitive. AGL21 positively regulates *ABI5* expression in seeds. Genetic analyses reveal that *AGL21* is epistatic to *ABI5* in controlling seed germination. Chromatin immunoprecipitation assays further demonstrate that AGL21 could directly bind to the *ABI5* promoter in plant cells. Taken together, our results suggest that AGL21 acts as a surveillance integrator that incorporates environmental cues and endogenous hormonal signals into ABA signaling to regulate seed germination and early post-germination growth.

## INTRODUCTION

Seed dormancy is a vital trait for plants to adapt to varieties of habitats and climates. Seed germination is arrested under adverse conditions and resumed when the conditions are favorable. Generally, seed germination commences with three phases of water uptake (Bewley, 1997), followed by embryo expansion and finished with radicle emergence. The optimal level of seed dormancy and germination has important repercussions on agricultural production. Therefore, it is essential to study the underlying molecular mechanisms that control seed dormancy and germination.

To sessile organisms, the ambient environment is vital for their survival. A variety of environmental factors, including nutrients, water availability, temperature, light, oxygen, and soil salinity, affect seed germination (Finkelstein et al., 2008; Holdsworth et al., 2008). Plants have evolved an array of strategies to constantly monitor the changing environmental conditions to decide when to germinate (Finkelstein et al., 2008). The environmental cues perceived by seeds can be incorporated into endogenous hormonal signaling pathways to regulate germination. Among various phytohormones, abscisic acid (ABA) and gibberellic acid (GA) are regarded as the primary regulators of transition from dormancy to germination (Seo et al., 2006; Shu et al., 2016). ABA is required for seed dormancy maintenance, while GA acts antagonistically to release dormancy and initiate seed germination (Shu et al., 2013). Environmental factors regulate the ABA:GA balance and the sensitivity to these hormones by modifying the biosynthetic and catabolic pathways, as well as signaling pathways, thus modulating seed dormancy and germination (Finch-Savage and Leubner-Metzger, 2006).

During germination, ABA content is decreased rapidly, and ABA signaling must be actively repressed. Through screening for ABA-insensitive (ABI) mutants, several ABA signaling components control seed germination have been identified in *Arabidopsis* (Finkelstein et al., 2002; Nambara and Marion-Poll, 2003). In contrast to ABI1/2, ABI3, ABI4 and ABI5 are key positive regulators of ABA signaling that modulate seed germination and post-germination development (Giraudat et al., 1992; Parcy et al., 1994; Finkelstein et al., 1998; Lopez-Molina et al., 2002). ABI5 is one of the 13 members of the group-A bZIP TF subfamily, and can directly bind to the ABA-RESPONSE ELEMENT (ABRE) *cis*-element in the promoter sequence of ABA-responsive genes, such as *Arabidopsis EARLY METHIONINE-LABELED 1* (*AtEm1*), *AtEm6*, and *RD29B* to modulate their expression (Carles et al., 2002; Finkelstein et al., 2005; Nakashima et al., 2006). ABI5 interacts with ABI3 and acts downstream of ABI3 to execute an ABA-dependent growth arrest during germination (Lopez-Molina et al., 2002). ABI5 protein level and activity are tightly regulated post-transcriptionally. In stress conditions, ABA triggers phosphorylation and activation of ABI5 (Lopez-Molina et al., 2001; Dai et al., 2013), and KEEP ON GOING (KEG) E3 ligase is rapidly degraded (Stone et al., 2006), promoting the accumulation of high levels of ABI5. In favorable conditions, ABI5 is dephosphorylated by FyPP/PP6 (for Phytochrome-associated serine/threonine protein phosphatase/Ser/Thr-specific phosphoprotein phosphatase 6), and then the inactive ABI5 is degraded by the 26S proteasome (Dai et al., 2013). ABI5 can also be modified by S-nitrosylation and sumoylation, and is rapidly degraded, which is facilitated by ABI FIVE BINDING PROTEIN (AFP) and KEG (Miura et al., 2009; Albertos et al., 2015).

The MADS-box gene family in higher plant is a large family with more than 100 members (De Bodt et al., 2005). These TFs are involved in almost every developmental process in plants (Smaczniak et al., 2012). However, their roles in seed germination are largely unexplored. Only two MADS-box genes were found regulating seed germination till now. One is *FLOWERING LOCUS C* (*FLC*)/*AGL25*, which is involved in temperature-dependent seed germination through influencing ABA catabolic pathway and GA biosynthetic pathway (Chiang et al., 2009). Another MADS-box gene *AGL67* may act as a repressor of seed germination, for knockout of it decreasing seed dormancy (Bassel et al., 2011). In our previous study, we found MADS-box TF AGL21 is involved in lateral root development and growth mediated by various environmental and physiological signals (Yu et al., 2014). In this study, we report that AGL21 also modulates seed germination by regulating ABA signaling. Overexpression of *AGL21* conferred hypersensitive seed germination to ABA, high NaCl and mannitol, while the *AGL21* knockout mutants showed the opposite phenotypes. Further analyses showed that *AGL21* was induced by various stresses, and highly expressed in dry seeds, but decreased quickly after imbibition and germination, which is coincident with that of *ABI5*. Genetic analysis showed that *AGL21* was involved in ABA signaling, acting downstream of *ABI1/2* and upstream of *ABI5*. Taken together, our results suggest that AGL21 integrate environmental signals and internal hormonal signals to ABA signaling by directly modulating *ABI5* to fine-tune seed germination and post-germination growth.

## RESULTS

### AGL21 negatively regulates seed germination and post-germination in response to ABA, salt and osmotic stress

*AGL21* was reported to be expressed in developing embryos (Burgeff et al., 2002), and our previous study showed that *AGL21* also had high expression levels in siliques and dry seeds besides in roots (Yu et al., 2014). These data indicates that AGL21 may play some roles in seed development or seed germination. To study the function of AGL21 in seed germination, we obtained 35S::*AGL21* (OX 1-6 and OX 3-5), 35S::*AGL21-HA* (OX 29-3) transgenic *Arabidopsis* lines, and two T-DNA insertion mutants: CS118325 (*agl21-1*) and GK_157C08 (*agl21-2*) from ABRC (Yu et al., 2014). Gene expression analyses by quantitative real-time polymerase chain reaction (qRT-PCR) showed that the expression of *AGL21* was abolished in the mutants while highly up-regulated in the *AGL21*-overexpressing plants (Fig. S1). To evaluate the function of AGL21 in seed germination, we germinated the seeds of *AGL21*-overexpressing plants, *agl21* mutants and wild-type (WT) plants on MS media with or without ABA. In the absence of ABA, the seed germination rates of different genotypes were similar (Fig. 1A, B). In the presence of different concentrations of ABA, *agl21-1* and *agl21-2* seeds were more resistant to ABA inhibition. On the contrary, *AGL21*-overexpressing seeds were more sensitive to ABA during germination (Fig. 1A-D). In line with the germination rate data, *agl21* mutant plants also showed higher cotyledon-greening percentages than WT plants, while the three *AGL21*-overxpressing lines exhibited significant lower cotyledon-greening rates after 8 days germination (Fig. 1E). These results suggested that AGL21 acts as a negative regulator of seed germination and post-germination.

**Figure 1.**
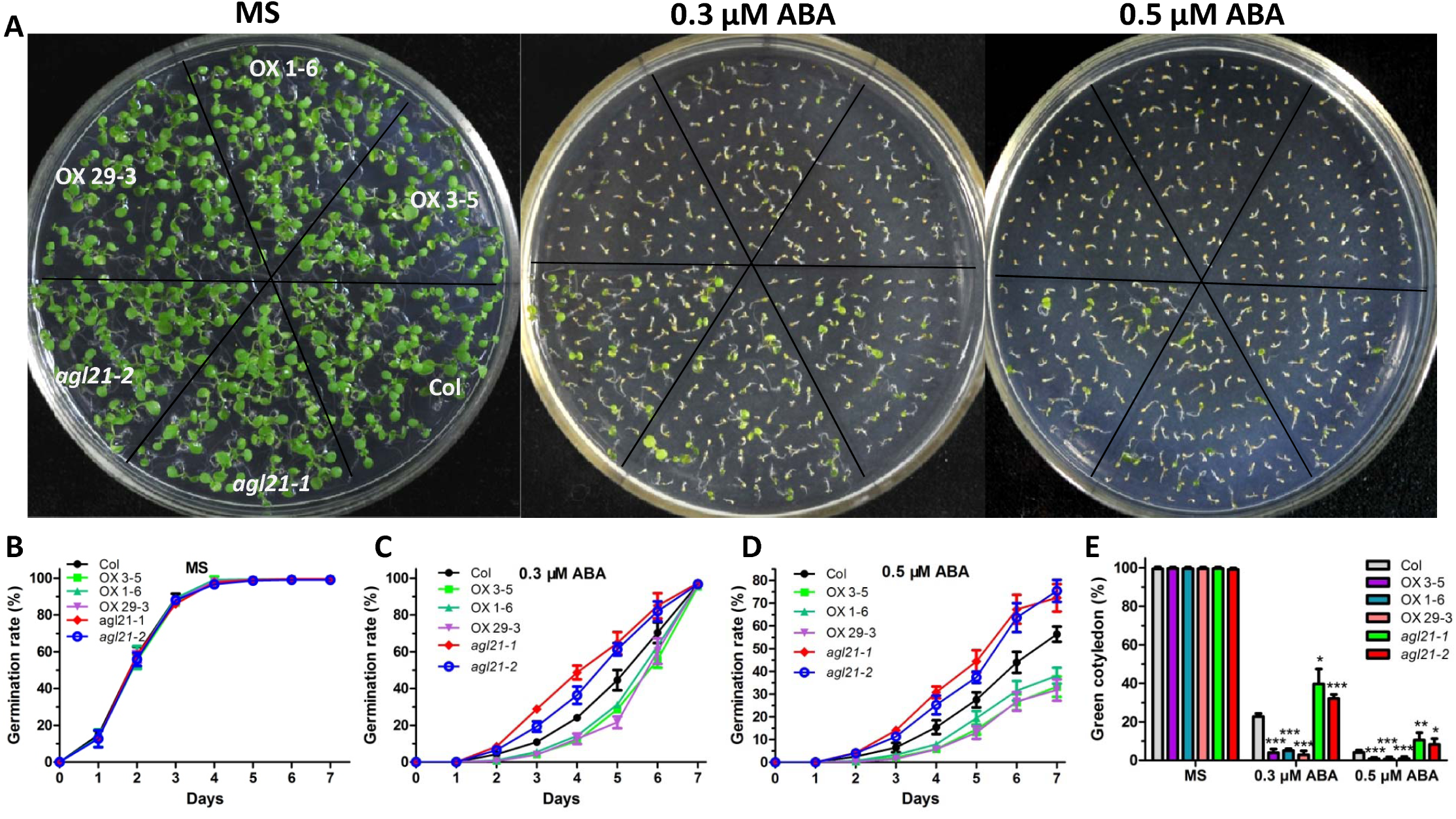
Response of *AGL21*-overexpressing and *agl21* mutant plants to NaCl and mannitol in seed germination. Seed germination assays were carried out as described in METHODS. Seed germination percentages of the indicated genotypes grown on MS medium or MS medium containing 150 mM NaCl or 300 mM mannitol were quantified every day from the 1st day to the 7th day after sowing. Cotyledon-greening percentages of the eighth day were recorded. Four independent experiments were conducted, with at least 36 seeds per genotype in each replicate. Values are mean ± SD of four replications. **(A)** Photographs of seedlings grown on different media at day 8 after the end of stratification. **(B-D)** Seed germination rates of indicated genotypes grown on different media. **(E)** Green cotyledon at day 8 after the end of stratification. Values are mean ± SD of four replications (*P<0.05,**P<0.01).

Since AGL21 modulates ABA-regulated seed germination, we tested whether AGL21 affects the seed germination response to salt stress. The seeds were sown on MS medium supplemented with 150 mM NaCl. Compared with WT seeds, germination and cotyledon-greening of *AGL21*-overexpressing seeds were more severely inhibited by NaCl, whereas the *agl21* mutant seeds showed much higher germination and cotyledon-greening ratios (Fig. 2A, C and E). To identify whether AGL21 is involved in salt-specific or general osmotic responses, we further tested seed germination on medium containing osmotic reagent mannitol. Similarly, on MS medium containing 300 mM mannitol, germination of *AGL21*-overexpressing seeds were much more sensitive while the *agl21* mutant seeds were insensitive to the inhibition effects of mannitol compared with WT seeds (Fig. 2A, D and E), indicating AGL21 negatively regulates seed germination in response to osmotic stress.

**Figure 2.**
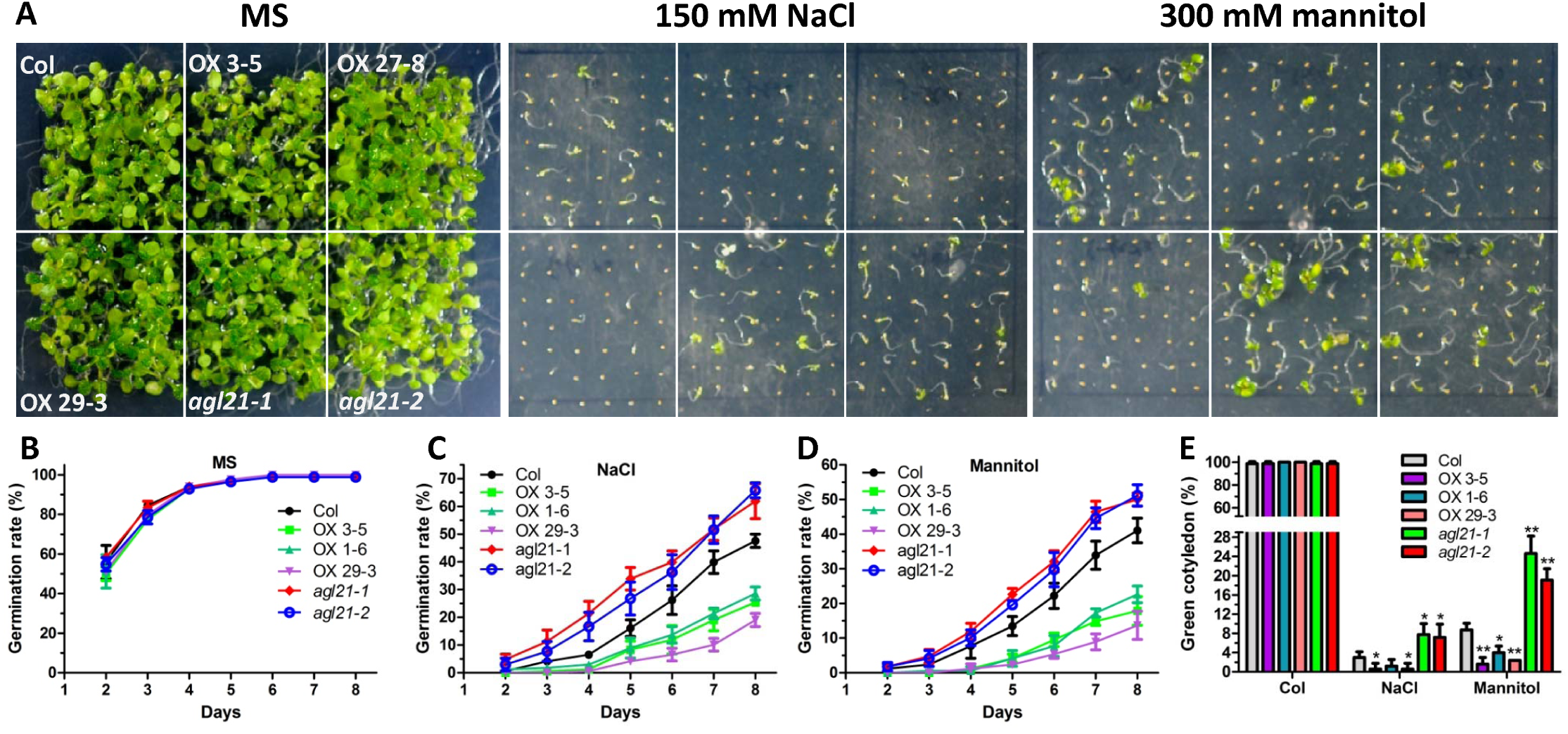
Response of *AGL21*-overexpressing and *agl21* mutant plants to NaCl and mannitol in seed germination. Seed germination assays were carried out as described in METHODS. Seed germination rates of the indicated genotypes grown on MS medium or MS medium containing 150 mM NaCl or 300 mM mannitol were quantified every day from the 2nd day to the 8th day after sowing. Cotyledon-greening percentages of the 8th day were recorded. Four independent experiments were conducted. At least 42 seeds per genotype were measured in each replicate. Values are mean ± SD of four replications. **(A)** Photographs of seedlings grown on different media at day 8 after the end of stratification. **(B-D)** Seed germination rates of indicated genotypes grown on different media. **(E)** Green cotyledon at day 8 after the end of stratification. Values are mean ± SD of four replications (*P<0.05,**P<0.01).

### *AGL21* is responsive to multiple stresses during seed germination

*AGL21* responds to multiple endogenous and exogenous signals, such as ABA, methyl jasmonate (MeJA), indole-3-acetic acid (IAA), nitrogen (N) and sulfur (S) starvation during root development (Yu et al., 2014). Our results in this study show that AGL21 is involved in the inhibition of germination in response to ABA, NaCl and osmotic stress. Thus we wanted to know whether *AGL21* also responded to these stress signals during germination stage. We analyzed the *AGL21* expression levels in seed germination stages on media with ABA, NaCl or mannitol. The results show that all these stress treatments significantly induced *AGL21* expression in germinating seed. Moreover, *AGL21* was also markedly induced by N deficiency, MeJA and IAA treatments during seed germination (Fig. 3A). These data support the surveillance function of AGL21 in seed germination in response to ABA, salt and osmotic stresses, and maybe other stresses.

**Figure 3.**
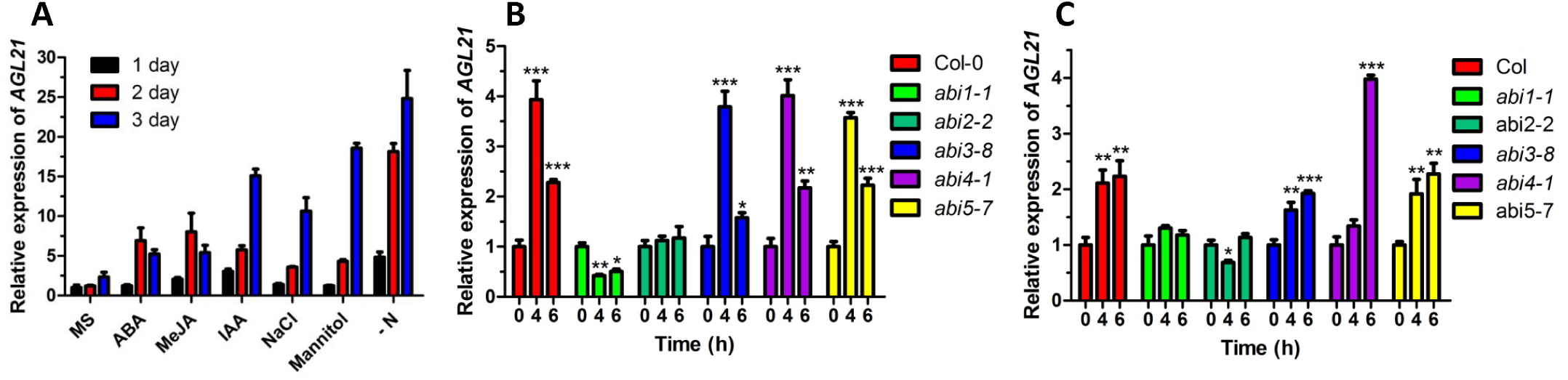
Response of *AGL21* to variety external signals in WT and *abi* mutants. **(A)** Response of *AGL21* to variety external signals in 2 day germinating seeds of WT. Seeds were stratified at 4℃ for 2 days in darkness and then sown on MS medium, MS medium without N (-N) or MS medium containing 100 mM NaCl, 250 mM mannitol, 0.2 μM ABA, 10 μM IAA, 10 μM MeJA, respectively. After 2 days germination, seeds were harvested for qRT–PCR analyses. **(B-C)** Response of *AGL21* to ABA (B) and mannitol (C) treatments in WT and different *abi* mutants. 4-day-old seedlings of different genotypes were transferred to MS solution containing 20 μM ABA or 350 mM mannitol, and harvested at indicated time points for RNA extraction and qRT–PCR analyses. The transcript levels of *AGL21* were normalized to the *UBQ5* expression. Values are mean ± SD of three replications.

### AGL21 is involved in ABA signaling

In order to study whether AGL21 is involved in ABA signaling, we checked the *AGL21* expression levels in *abi* mutants in response to ABA. As shown in Figure 3B and 3C, ABA and mannitol treatments could significantly induce *AGL21* expression in *abi3-8, abi4-1, abi5-7* background plants as that in WT background plants. However, in *abi1-1* and *abi2-2* background plants, no significant induction of *AGL21* transcription was observed. These data suggested that AGL21 might function in ABA signaling pathway, and may act downstream of ABI1 and ABI2, and upstream or in parallel with ABI3, ABI4 and ABI5.

We further analyzed the expression levels of several ABA signal pathway genes in germinating seeds of *AGL21*-overexpressing, knockout and WT plants germinated on MS medium or MS medium with ABA. We found that AGL21 positively regulated some downstream ABA signal pathway genes, such as *ABI5, AtEM6, RD29B* and *ABA RESPONSE ELEMENT-BINDING FACTOR 2 (AREB2)/AREB BIND FACTOR 4* (*AREB2*/*ABF4*) (Fig. 4A-H). However, gene expression levels of upstream ABA signal pathway genes, such as *ABI1, ABI2, SNF1-RELATED PROTEIN KINASE 2.2* (*SnRK2.2*) and *SnRK2.3*, did not change significantly in *AGL21*-overexpressing, knockout and WT plants (Fig. 4I-P). Together, these results indicate that AGL21 is involved in ABA signaling, and may act downstream of TYPE 2C PROTEIN PHOSPHATASES (PP2Cs) and SnRKs and upstream of ABI5 and other downstream ABA signal components to regulate seed germination and post-germination growth.

**Figure 4.**
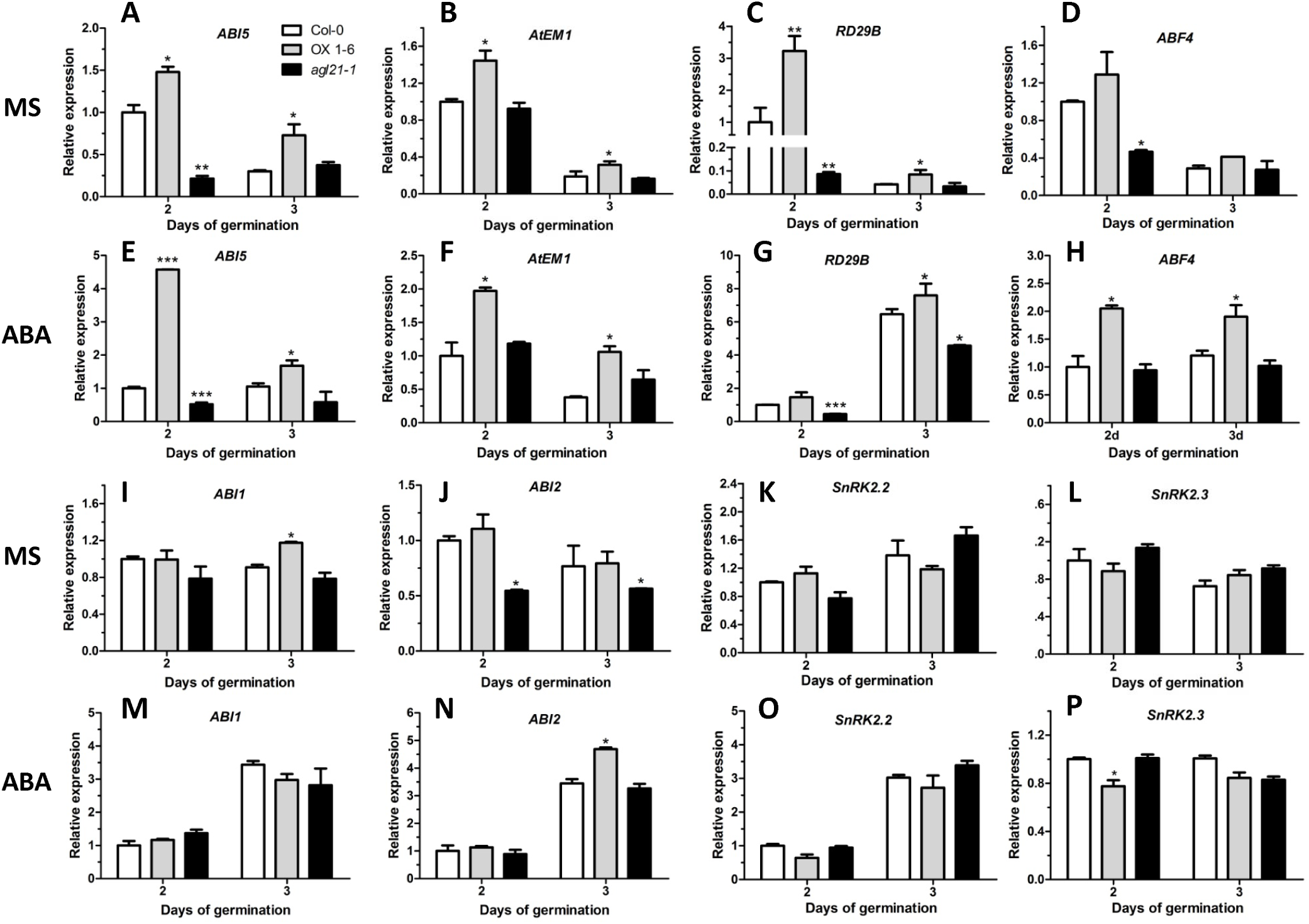
Expression levels of ABA signal pathway genes in *agl21-1* mutant and *AGL21*-overexpressing lines quantified by qRT-PCR. Values are mean ± SD of three replications (*P< 0.05, **P< 0.01, ***P< 0.001). *UBQ5* was used as an internal reference. **(A-D)** Expression levels of *ABI5, AtEM6, RD29B, ABF4* in 2-day-old and 3-day-old *AGL21*-overexpressing and *agl21-1* plants grown on MS medium. **(E-H)** Expression levels of *ABI5, AtEM6, RD29B, ABF4* in 2-day-old and 3-day-old *AGL21*-overexpressing and *agl21-1* plants grown on MS medium containing 0.15 μM ABA. **(I-L)** Expression levels of *ABI1, ABI2, SnRK2.2, SnRK2.3* in 2-day-old and 3-day-old *AGL21*-overexpressing and *agl21-1* plants grown on MS medium. **(M-P)** Expression levels of *ABI1, ABI2, SnRK2.2, SnRK2.3* in 2-day-old and 3-day-old *AGL21*-overexpressing and *agl21-1* plants grown on MS medium containing 0.15 μM ABA.

### AGL21 does not affect ABA and GA biosynthesis

ABA and GA antagonistically govern seed germination. ABA promotes seed dormancy, while GA stimulates seed germination. To investigate whether AGL21 regulates seed germination by altering ABA or GA content, we analyzed the expression levels of many key genes in ABA biosynthesis pathway, such as *ABA DEFICIENT 2* (*ABA2*), *ABA3, ARABIDOPSIS ALDEHYDE OXIDASE 3* (*AAO3*), *9-CIS-EPOXYCAROTENOID DIOXYGENASE* (*NCED3*), and key ABA catabolic enzyme family genes *CYP707A1*–*CYP707A4*. Our data show that all of these genes had similar expression levels in day 2 and day 3 germinating seeds of different genotypes both on MS and ABA media (Fig. S2). Moreover, we tested the ABA contents in dry seeds, and found there was no significant change in ABA contents between the WT, *agl21-1* and *AGL21*-overexpressing seeds (Fig. S3).

We also analyzed the expression levels of several key genes in GA biosynthesis pathway, including *ENT-COPALYL DIPHOSPHATE* (*CPS*), *ENT-KAURENE SYNTHASE* (*KS*), *ENT-KAURENOIC ACID OXIDASE 2* (*KAO2*), *GA20-OXIDASE 1* (*GA20OX1*), *GA20OX2, GA20OX3, GA REQUIRING 3* (*GA3*), *GA3-OXIDASE 1* (*GA3OX1*), *GA3OX2* and *GA3OX3*, in day 3 germinating seeds. qRT-PCR analysis revealed that the expression levels of the examined GA metabolism genes were largely unaltered in the *AGL21*-overexpressing, knockout and WT seeds grown on MS medium with or without ABA (Fig. S4). Taken together, these results show that AGL21 does not alter ABA or GA content during seed germination and post-germination growth.

### *AGL21* has similar expression pattern to *ABI5* and regulates *ABI5* expression and protein accumulation in seeds

To uncover the molecular networks underlying AGL21 regulation of seed germination, we monitored the expression levels of genes associated with ABA dependent seed germination, such as *ABI1, ABI2, ABI3, ABI4, ABI5, MYB DOMAIN PROTEIN 96* (*MYB96*), *ACYL-COENZYME A-BINDING PROTEIN 1* (*ACBP1*), *ARABIDOPSIS HISTIDINE KINASE 1* (*AHK1*), *RING-H2 FINGER A2A* (*RHA2a*), *RELATED TO ABI3/VP1 1* (*RAV1*) (Tran et al., 2007; Bu et al., 2009; Du et al., 2013; Feng et al., 2014; Lee et al., 2015), in day 3 germinating seeds in the present of ABA. Among these genes, *ABI3, ABI4, ABI5* were found significantly up-regulated in *AGL21*-overexpressing plants. However, their expression was not significantly down-regulated in *agl21-1*, except *ABI5* (Fig. 5A), indicating that *ABI5* might be a target of *AGL21*.

**Figure 5.**
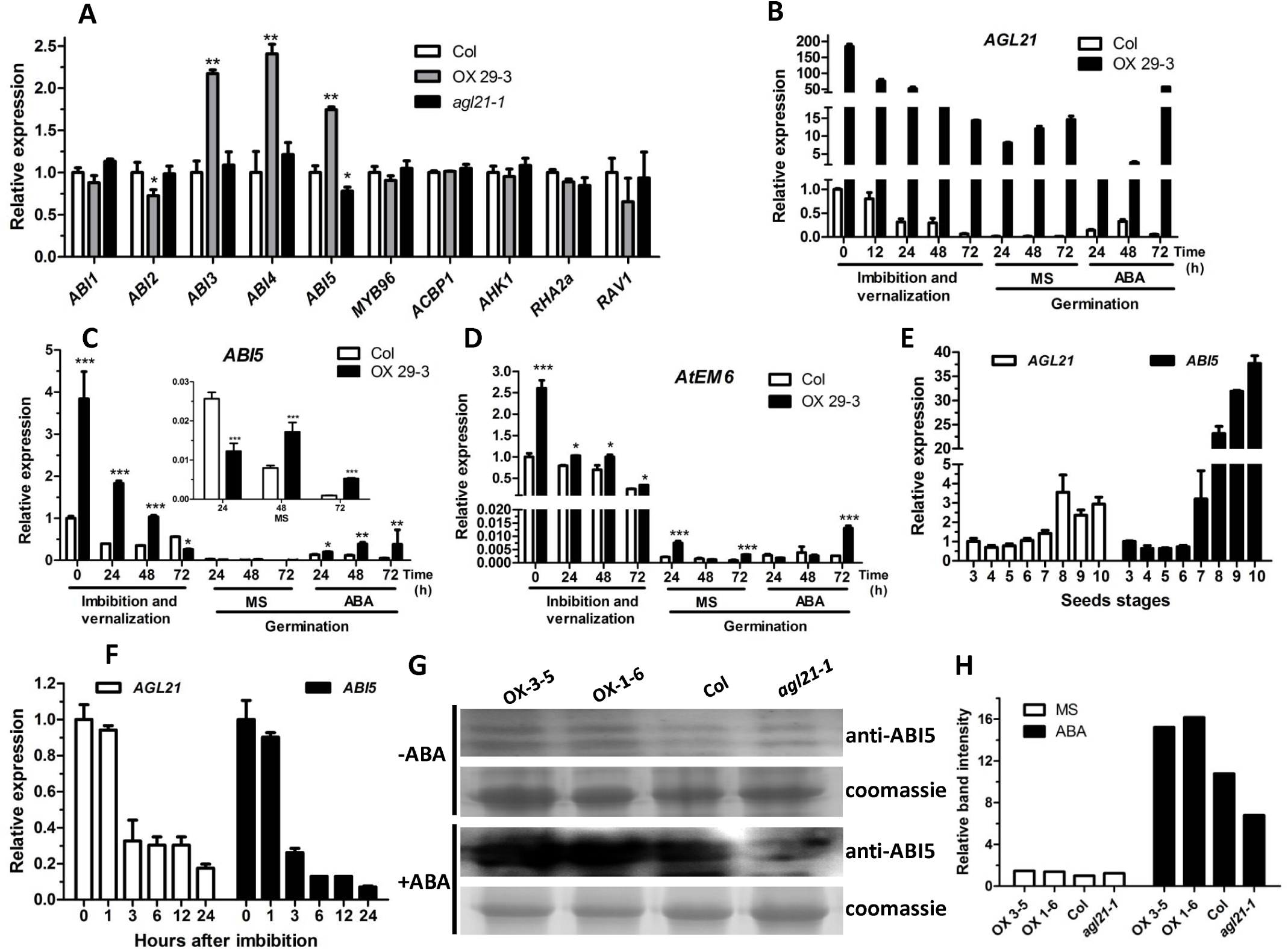
*AGL21* has similar expression pattern to *ABI5* and positively regulates *ABI5* expression and protein accumulation. **(A)** Expression levels of genes involved in ABA-dependent seed germination in WT, *AGL21*-overexpressing and *agl21-1* germinating seeds. Seeds sown on MS medium supplemented with 0.2 μM ABA for 3 days, and then harvested for RNA extraction and qRT-PCR analyses. *UBQ5 was* used as an internal reference. Values are mean ± SD of three replications (*P< 0.05, **P< 0.01). **(B-D)** *AGL21, ABI5* and *AtEM6* expression patterns in imbibed seeds and germinating seeds on MS medium or MS medium containing 0.15μM ABA. Seeds were imbibed in water at 4 E or sown on medium containing ABA or not after 2 days imbibition at 4 E, and harvested at indicated time points for RNA extraction and qRT-PCR analyses. *UBQ5* was used as an internal reference. Values are mean ± SD of three replications (*P< 0.05, **P< 0.01, ***P< 0.001). **(E-F)** Expression of *AGL21* and *ABI5* during seed maturation and imbibition. Expression data were extracted from the *Arabidopsis* eFP browser. The eFP browser was set to the Developmental Map and the Seed, with absolute values for gene expression. **(G-H)** ABI5 protein levels in WT, *AGL21*-overexpressing and *agl21-1* germinating seeds. 3 days germinating seeds sown on MS medium or MS medium containing 0.15 uM ABA were used for Western blot analysis with anti-ABI5 antibody. A nonspecific coomassie blue-stained band is shown as a loading control. Relative band intensity was measured using ImageJ software (NIH).

Moreover, we found high levels of *AGL21* transcripts accumulated in WT dry seeds, but the levels gradually dropped after 12-72 hours imbibition and continually decreased to a very low level after 1-3 days germination (Fig. 5B). However, when germinated on medium with ABA, *AGL21* expression was significantly induced by ABA during germination. Similar expression pattern of *AGL21* in *AGL21*-overexpressing seeds was also observed, except small increases during 1-3 days germination on MS medium (Fig. 5B), which were probably due to the higher expression of *AGL21* in developing roots. The pattern of *AGL21* expression is similar to that of *ABI5* (Okamoto et al., 2010), and coincides with the content changes of ABA, which is also high in dry seeds but reduces rapidly after imbibition (Ali-Rachedi et al., 2004). We then compared the expression levels of *ABI5* in dry seeds, imbibed seeds and germinating seeds of *AGL21*-overexpressing and WT plants (Fig. 5C). *ABI5* showed a very similar expression pattern to *AGL21*. More importantly, expression of *ABI5* was significantly higher in *AGL21*-overexpressing seeds compared with that in WT seeds in most time-points checked, especially in seeds imbibed for 0-48 h and seeds germination on ABA medium for 24-72 h (Fig. 5C). In addition, one of ABI5 target genes, *AtEM6*, also had a similar expression pattern to *ABI5* and *AGL21*, and also had increased expression levels in *AGL21*-overexpressing imbibed seeds, seeds germination on MS medium for 24 h and 72 h and seeds grown on ABA medium for 72 h (Fig. 5D). Furthermore, we also compared the expression profiles of *AGL21* and *ABI5* in seed using public expression data extracted from the *Arabidopsis* eFP browser (Winter et al., 2007). As data showed in Figure 5E, *AGL21* and *ABI5* have similar expression patterns in different seed development stages, with much higher expression levels of both genes at the late stages of seed development. Both of these two genes have the highest expression levels in dry seeds and dramatically decrease after 3-24 h imbibition (Fig. 5F). These data are consistent with our qRT-PCR analyses, implying that AGL21 may directly regulate *ABI5* expression in seeds.

Since ABI5 protein contents directly affect ABA-dependent seed germination and post-germination growth, we examined ABI5 protein levels in seeds of WT, *AGL21*-overexpressing and *agl21-1* in the present or absence of exogenous ABA. In seeds germinated on MS medium for 3 days, no clear differences of ABI5 protein level were detected between the seed of different genotypes. However, in the present of ABA, significantly higher ABI5 protein levels accumulated in *AGL21*-overexpressing seeds, while significantly decreased in *agl21-1* seeds compared with WT seeds (Fig. 5G and H). All these results suggest that AGL21 modulates ABI5 at both the transcriptional and protein levels.

### *ABI5* is one of the target genes of AGL21 in regulating ABA-dependent seed germination

The expression pattern *of AGL21* shows high similarity to that of *ABI5* in seeds and its function in seed germination also similar to *ABI5*. To test whether AGL21 acts in the same pathway to ABI5 in seed germination, we generated 35S::*AGL21* plants in the genetic background of the *abi5-7* mutant (*abi5-7*/*AGL21*OX 1-6). As shown in ABA response assays, *AGL21*OX 1-6 seeds were hypersensitive, however, the *abi5-7*/ *AGL21*OX 1-6 plants showed an ABA insensitive phenotype as that of *abi5-7* mutant (Fig. 6), suggesting that AGL21 act upstream of ABI5 in ABA signaling pathway to regulate seed germination.

**Figure 6.**
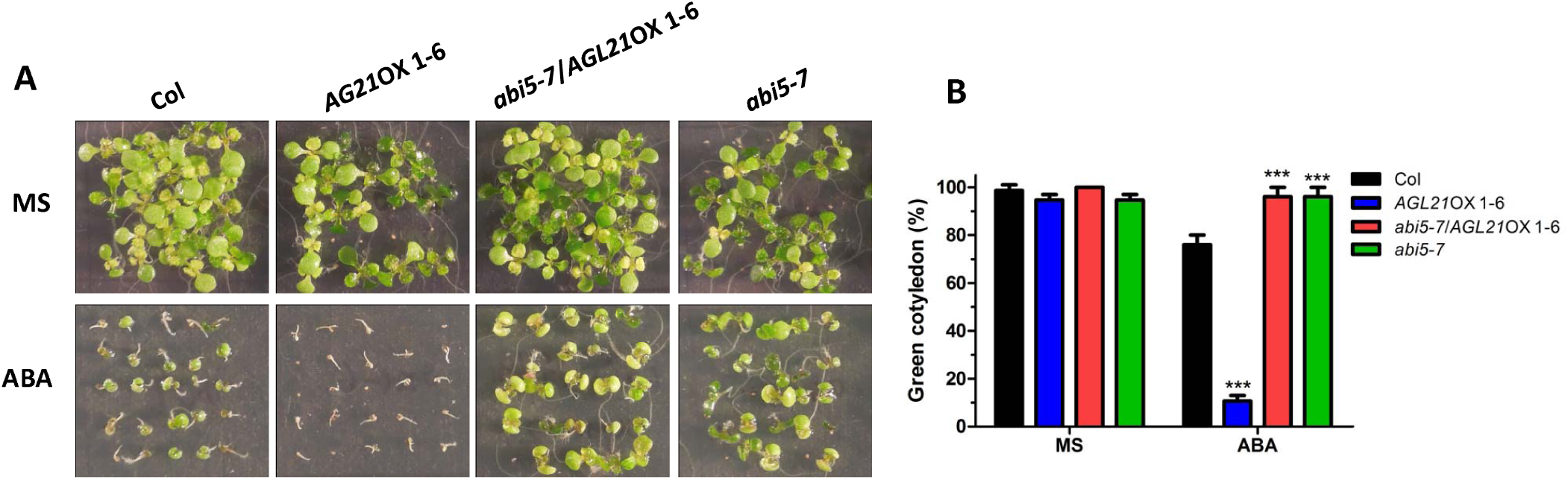
Genetic relationship between *AGL21* and *ABI5.* **(A)** ABA response of WT, *abi5-7, AGL21*-overepressing plants and *AGL21*-overepressing plants in the genetic background of *abi5-7 (<u>abi5-7</u>*<u>/</u>*AGL21* OX 1-6). Seeds of different genotypes were stratified at 4 ᎚ for 2 days and then sown on MS medium or MS medium containing 0.8 uM ABA for 12 days. **(B)** Green cotyledon percentage of seeds grown on MS medium or MS medium containing 0.8 uM ABA for 12 days. Values are mean ± SD of three replications. At least 25 seeds per genotype were counted in each replicate (***P< 0.001).

MADS-box TFs can modulate their target genes expression by specifically binding as homo- or heterodimers to the flexible CArG-box (C-[A/T]rich-G) *cis*-element (Riechmann et al., 1996). Sequence analysis found that the *ABI5* promoter contains six putative binding sites (Fig. 7A), which contained a core sequence that meets the patterns for any of the possible three CArG-box motifs, C(A/T)_8_G, C(C/T)(A/T)_6_(A/G)G, or C(C/T)(A/T)G(A/T)_4_(A/G)G (de Folter and Angenent, 2006; Ito et al., 2008; Fujisawa et al., 2011). The presence of CArG-box motifs led us to examine whether AGL21 is targeted to the *ABI5* promoter.

**Figure 7.**
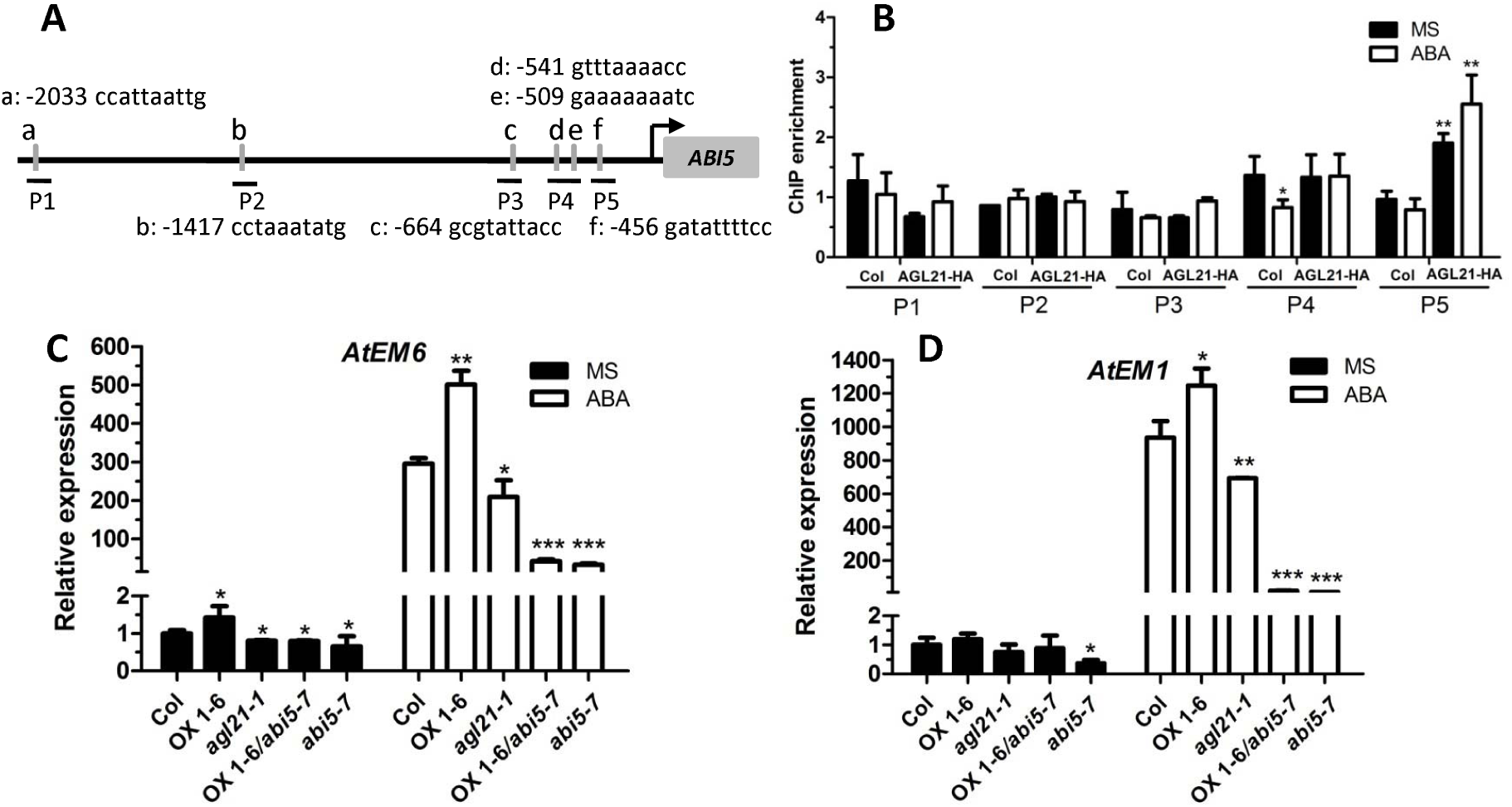
AGL21 directly regulates *ABI5* expression. **(A)** Schematic representation of *ABI5* promoter showing putative CArG-box motifs upstream of the transcription start site. CArG-box motifs are indicated with gray lines, above/below which the sequence and the sites of the last base of the motif relative to the start code are shown. PCR-amplified fragments are indicated by black lines under the CArG-box motifs. **(B)** ChIP-qPCR assay of AGL21 binding to *ABI5* promoter. The 5-day-old 35S::AGL21-HA (OX 29-3) transgenic plants and WT plants were transferred to MS solution with or without 10 μM ABA for 4 h, and then the seedlings were harvested for ChIP-qPCR assay using anti-HA antibody. Enriched values were normalized with the level of input DNA. Values are mean ± SD of three replications (*P< 0.05, **P< 0.01). **(C)** Expression of *AtEM1* and *AtEM6* in WT, *agl21-1, abi5-7, AGL21*-overepressing plants and *AGL21*-overepressing plants in the genetic background of *abi5-7* (*abi5-7/AGL21OX* 1-6). Seeds of different genotypes were stratified at 4 ᎚ for 2 days before sown on MS medium or MS medium containing 0.2 μM ABA for 3 days, and then were harvested for qRT-PCR analyses. *UBQ5 was* used as an internal reference. Values are mean ± SD of three replications (*P< 0.05, **P< 0.01, ***P< 0.001).

We performed chromatin immunoprecipitation (ChIP) assays using 4-day-old 35S::*AGL21-HA* transgenic plants grown on MS medium or MS medium containing 0.15 μM ABA. DNA fragments bound to epitope-tagged AGL21 proteins were analyzed by qPCR assays. The ChIP-qPCR analysis demonstrated that the P5 DNA fragment was significantly enriched by AGL21 both in plants grown on MS medium or MS medium containing ABA, with much higher enrichment in the plants grown in the present of ABA (Fig. 7B). However, other four genomic fragments containing putative CArG-box motif were not enriched. In addition, none of the five genomic fragments was found enriched by AGL21 in WT plants (Fig. 7B). All these data support the specific interaction of AGL21 with the P5 region of the *ABI5* promoter.

To verify that AGL21 directly regulates *ABI5* expression, we checked the expression levels of ABI5-targeted genes *AtEM1* and *AtEM6* in seeds sown on MS media or MS media with 0.2 μM ABA for three days. As shown in Figure 7C, on MS medium, *AtEM6* was up-regulated in *AGL21*-overexpressing seeds, and significantly down-regulated in *agl21-1, AGL21OX* 1-6/*abi5-7*, and *abi5-7* compared with WT. As for *AtEM1*, only significantly decreased expression level in *abi5-7* seeds was observed on MS medium (Fig. 7D). On MS medium supplemented with ABA, the expression levels of both *AtEM1* and *AtEM6* markedly increased in *AGL21*-overexpressing seeds, while significantly reduced in *agl21-1* seeds. However, when *AGL21* was overexpressed in *abi5* background (*AGL21*OX 1-6/*abi5-7*), similar expression levels of *AtEM1* and *AtEM6* were observed to that of *abi5-7* seeds (Fig. 7C and D), indicating that AGL21-upregulated *AtEM1* and *AtEM6* expression is *ABI5*-dependent. These results confirm that *ABI5* is a target gene of AGL21.

## DISCUSTION

There are more than 100 MADS-box genes found in *Arabidopsis* genome (Gramzow and Theissen, 2013), with functions in the morphogenesis of almost all plant organs and throughout the whole life cycle (Smaczniak et al., 2012). However, their functions in seed dormancy and germination are largely unknown. So far, only two MADS-box gene, *FLC* and *AGL67*, were reported involved in seed germination (Chiang et al., 2009; Bassel et al., 2011). Recently, gene expression profiling analysis identified three MADS-box genes, including *AGL21*, were differentially expressed between imbibed dormant and after-ripened ecotype C24 seeds, suggesting putative functions of these genes in seed dormancy and germination (Barrero et al., 2010). In this study, we discovered that AGL21 regulated ABA-mediated seed germination and post-germination growth by modulating ABA signaling.

We found that AGL21 acts as a negative regulator in seed germination. *AGL21* is primarily expressed in root, but also has high expression levels in siliques and dry seeds, and it responds to multiple environmental and internal signals both in germinating seeds and roots (Fig. 3) (Yu et al., 2014). Overexpression of *AGL21* results in hypersensitivity of seed germination to ABA, high salt, and osmotic stress, while knockout *AGL21* confers opposite phenotypes during germination (Fig. 1 and Fig. 2). Seed germination is tightly regulated by ABA:GA balance and ABA signaling in seed (Finch-Savage and Leubner-Metzger, 2006). We found that AGL21 does not affect ABA and GA biosynthesis or catabolism (Fig. S2-S4), indicating that AGL21-regulated seed germination is not through affecting ABA or GA content. Our further analyses of *AGL21* expression in *abi* mutants (Fig. 4) and ABA signal pathway genes expression in *agl21-1, AGL21*-overexpression, and WT seeds (Fig. 5) indicate that *AGL21* is involved in ABA signal pathway to regulate seed germination and may acts downstream of PP2Cs and SnRKs, but upstream of *ABI5* and other ABA-responsive genes.

As a key player in ABA-triggered arrest of germination and post-germination growth, ABI5 is regulated at both transcriptional and post-transcriptional levels. Several genes, such as *WRKY2, RAV1, MYB7, SALT- AND DROUGHT-INDUCED RING FINGER1* (*SDIR1*), *SDIR1-INTERACTING PROTEIN1* (*SDIRIP1*), *HY5, B-BOX21* (*BBX21*), *DELAY OF GERMINATION 1* (*DOG1*), NUCLEAR FACTOR-Y C- RGA-LIKE 2 (NF-YC–RGL2), were reported to regulate seed germination by modulating *ABI5* expression directly or indirectly (Zhang et al., 2007; Chen et al., 2008; Jiang and Yu, 2009; Feng et al., 2014; Xu et al., 2014; Kim et al., 2015; Zhang et al., 2015; Dekkers et al., 2016; Liu et al., 2016). Our current study implicates that AGL21 can directly regulate *ABI5* expression during seed germination. Firstly, we found *AGL21* and *ABI5* had similar expression patterns during seed development and germination. *ABI5* is expressed throughout seed development, reaching the highest transcript level at mature seed stage, but dropping during imbibition and germination unless exposed to stresses (Fig. 5C) (Brocard et al., 2002). Here, we found that the expression pattern of *AGL21* in seeds is in accord with that of *ABI5* (Fig. 5B- 5F) and ABA content changes in seeds during the same period. These results imply a possible regulation between these two genes. Further qRT-PCR analyses found that among 10 seed germination related genes, only *ABI5* had significantly increased expression level in *AGL21*-overexpressing germinating seeds, with significantly reduced expression in *agl21-1* seeds (Fig. 5A). Then we found *ABI5* and its target gene *AtEM6* had markedly elevated expression in *AGL21*-overexpressing dry seeds, imbibed seeds, germinating seeds grown on MS or ABA media (Fig. 5C and D). Moreover, more ABI5 protein was accumulated in *AGL21*-ovexpression germinating seeds, while down-regulated in *agl21-1* seeds on ABA media at the same time (Fig. 5G and H). In addition, genetics evidence found that AGL21-regulated seed germination is depend on ABI5. When overexpressing *AGL21* in *abi5* background (*abi5-7*/AGL21OX1-6), the hypersensitive seed germination to ABA was abolished, instead *abi5-7*/AGL21OX1-6 seeds showed similar sensitivity to *abi5* (Fig. 6). Taken together, these data suggest that AGL21 acts upstream of ABI5, and may directly modulate *ABI5* transcription.

To verify whether *ABI5* is the direct target of AGL21, ChIP-qPCR assay was carried out. We found one promoter DNA fragment containing a putative CArG-box *cis*-element of *ABI5* was significantly enriched in 4-day-old plants grown on MS or MS medium containing ABA (Fig. 7A and B), indicating that AGL21 directly binding to *ABI5* promoter to regulate its expression. *AtEM1* and *AtEM6*, two target gene of ABI5 (Carles et al., 2002), were markedly up-regulated in *AGL21*-overexpressing plants and down-regulated in *agl21-1* plants as that of *ABI5*. However, once ABI5 was mutated in *AGL21*-overexpressing background, the induction of *AtEM1* and *AtEM6* expression was blocked (Fig. 7C and D). Together, these data support the idea that AGL21 regulates seed germination by directly modulating *ABI5*. Although we also found other seed germination regulator genes, such as *ABI3, ABI4* and *ABF4*, were up-regulated in *AGL21*-overexpressing plants (Fig. 5H and Fig. 5A), the relationships of these genes to AGL21 await further investigation.

Timing of germination is a complex biological process that is regulated through intricate signaling pathways integrating diverse environmental signals, such as light, temperature, water, soil salinity and nutrition, into internal developmental programs, such as endogenous hormone signaling (Finkelstein et al., 2008; Jiang et al., 2016). For example, both osmotic and salinity stresses inhibit seed germination by affecting ABA signal pathway (Llanes et al., 2016). NO^3−^ can act as a seed germination enhancer by decreasing the level of ABA in the seed (Finkelstein et al., 2008; Osuna et al., 2015; Yan et al., 2016). Sulfate availability affects germination response to ABA and salt stress in *Arabidopsis* by regulating ABA biosynthesis (Cao et al., 2014). GA, ethylene, brassinosteroids and cytokinin have been shown to promote seed germination, while ABA, auxin, JA, SA inhibit seed germination (Kucera et al., 2005; Finkelstein et al., 2008; Wang et al., 2011; Liu et al., 2013; Shu et al., 2016). All these different phytohormones modulate seed germination most likely by regulating the ABA/GA balance at either the signaling or biogenesis levels, with ABI5 as one of the pivots involved in hormone crosstalk (Shu et al., 2016). Thus, the molecular links that incorporate external signals into internal plant hormonal signaling are crucial for seeds to germinate at proper time in the changing environment. We have demonstrated in this paper that AGL21 is a negative regulator of seed germination. *AGL21* responds to a variety of internal and external signals, such as ABA, MeJA, IAA, osmotic stress, salt stress, N and S deficiency (Fig. 3A), which are known to affect seed germination. Therefore, we propose that AGL21 may serve as environmental surveillance for seed germination. It controls seed germination by regulating ABI5 according to its environmental surveillance, which prevents seed germination under adverse conditions, an adaptive mechanism for plant survival.

## METHODS

### Plant Material and Growth Conditions

*Arabidopsis* Columbia-0 (Col-0) ecotype was used in this study. *AGL21*-overexpressing lines, such as *AGL21*OX 1-6 and *AGL21*OX 3-5, and mutants, such as*agl21-1* and *agl21-2* were reported previously (Yu et al., 2014). 35S::*AGL21-HA* transgenic line *AGL21OX* 29-3 was obtained from *Agrobacterium tumefaciens-mediated* transformation with *35S:AGL21-HA* construct. To get *35S::AGL21-HA* construct, the coding region of *AGL21* was amplified and cloned into pDONR207 with the primers *AGL21-HA* LP and *AGL21-HA* RP, and then shuttled it into pCB2004 vector (Lei et al, 2007). The *AGL21OX <u>1-6/abi5-7</u>* double mutant was generated by genetic cross of *AGL21OX* 1-6 and *abi5-7* mutant. All plants were grown at 220 under long-day condition (16-h light/8-h dark cycles).

### Seed germination assays

Seeds were collected at the same time were used for germination assays. Harvested seeds were aired dried at room temperature at least 3 weeks before the germination assays. For seed germination assays, seeds of each genotype were surface sterilized with 10% bleach for 12 min and washed five times with sterile water, and then were stratified at 40 for 2 days in darkness before sown on MS medium (1% sucrose, 0.5 % agar, pH 5.8) or MS medium supplemented with ABA, NaCl or mannitol. Seeds were germinated at 220 under 16-h light/8-h dark cycles. Germination (emergence of radicles) and post-germination growth (green cotyledon appearance) were scored at the indicated time points.

### qRT-PCR assay

Total RNA of seedlings was extracted with Trizol reagent (Invitrogen) and RNAs from dry seeds, imbibed seeds and germinating seeds were isolated using a TRIzol-based two-step method as previously described (Meng and Feldman, 2010). Total RNA samples were pretreated with DNase I (RNase Free) and 1.5 μg of total RNA was used for reverse transcription with oligo (dT)1_8_ to synthesize first-strand cDNA. qRT-PCR was performed with a StepOne Plus Real Time PCR System by using a TaKaRa SYBR Premix Ex Taq II reagent kit as described previously (Yu et al, 2013). All primers used are listed in Supplemental Table S1.

### Quantification of ABA

Dry seeds were ground in liquid nitrogen and ABA contents were measured by the ABA immunoassay kit as described (Yang et al, 2001).

### ChIP-qPCR assay

The ChIP assay was performed as reported previously (Cai et al., 2014). 35S::*AGL21-HA* transgenic plants (OX 29-3), anti-HA antibodies (Abmart), and salmon sperm DNA/protein A agarose beads (Millipore, USA) were used for ChIP experiment. DNA was purified using phenol/ chloroform (1:1, v/v) and precipitated. The enrichments of DNA fragments was quantified by qPCR using specific primers (Supplemental Table S1). Enriched values were normalized with the level of input DNA.

### Western blot

For western blot analysis, germinating seeds were powdered with liquid nitrogen and proteins were extracted with RIPA buffer [50 mmol/L Tris-HCl, pH 8.0, 0.1% Nonidet P-40,150 mmol/L NaCl, 1% sodium dodecyl sulfate, 0.5% sodium deoxycholate, and protease inhibitor cocktail tablets (Roche)]. Protein contents were determined using the Bradford method. Proteins were separated on 12% SDS-PAGE and electrotransferred to nitrocellulose membrane (Immobilon-P, MILLIPORE Corporation, USA). Anti-ABI5 antibody (Abiocode) at 1:1000 dilution was used for protein immunoblotting as previously described (Liu and Stone, 2010). The results were detected using a CCD camera system (Image Quant LAS 4000) using Super Signal West Femto Trial Kit (Thermo, USA).

### Statistical analysis

Statistically significant differences were computed based on the Student’s *t*-tests.

## SUPPLEMENTARY DATA

**Figure S1.** Expression level analyses of *AGL21*-overexpressing and knockout mutants.

**Figure S2.** Expression levels of ABA biosynthetic and catabolic pathway genes.

**Figure S3.** ABA contents in dry seeds of WT, *AGL21*-overexpressing and knockout mutant plants.

**Figure S4.** Expression levels of GA biosynthetic pathway genes.

**Table S1.** Primers used for PCR.

## FUNDING

This work was supported by the China National Natural Science Funds for Distinguished Young Scholar (grant no. 31500231), China Postdoctoral Science Foundation, No.9 Special Fund (grant no.2016T90577).

## AUTHOR CONTRIBUTIONS

L.-H.Y. and C.-B.X. designed the experiments; L.-H.Y. performed experiments and data analysis, and wrote the manuscript; J.W., Z.-Q.M., P.-X.Z., and Z.W. contributed to assist in performing part of the experiments; C.-B.X. supervised the project and revised the manuscript.

## ACKNOWLEDGEMENTS

We thank Dr. Chuan-You Li (Institute of Genetics and Developmental Biology, Chinese Academy of Sciences) for kindly providing *abi5-7* mutant. We also thank the ABRC for providing T-DNA insertion lines used in this study.

## REFERENCES

Albertos, P., Romero-Puertas, M.C., Tatematsu, K., Mateos, I., Sanchez-Vicente, I., Nambara, E., and Lorenzo, O. (2015). S-nitrosylation triggers ABI5 degradation to promote seed germination and seedling growth. Nat Commun 6, 8669, doi: 10.1038/ncomms9669.

Ali-Rachedi, S., Bouinot, D., Wagner, M.H., Bonnet, M., Sotta, B., Grappin, P., and Jullien, M. (2004). Changes in endogenous abscisic acid levels during dormancy release and maintenance of mature seeds: studies with the Cape Verde Islands ecotype, the dormant model of Arabidopsis thaliana. Planta 219, 479–488.

Barrero, J.M., Millar, A.A., Griffiths, J., Czechowski, T., Scheible, W.R., Udvardi, M., Reid, J.B., Ross, J.J., Jacobsen, J.V., and Gubler, F. (2010). Gene expression profiling identifies two regulatory genes controlling dormancy and ABA sensitivity in Arabidopsis seeds. Plant J 61, 611–622.

Bassel, G.W., Lan, H., Glaab, E., Gibbs, D.J., Gerjets, T., Krasnogor, N., Bonner, A.J., Holdsworth, M.J., and Provart, N.J. (2011). Genome-wide network model capturing seed germination reveals coordinated regulation of plant cellular phase transitions. Proc Natl Acad Sci U S A 108, 9709–9714.

Bewley, J.D. (1997). Seed Germination and Dormancy. Plant Cell 9, 1055–1066.

Brocard, I.M., Lynch, T.J., and Finkelstein, R.R. (2002). Regulation and role of the Arabidopsis abscisic acid-insensitive 5 gene in abscisic acid, sugar, and stress response. Plant Physiol 129, 1533–1543.

Bu, Q., Li, H., Zhao, Q., Jiang, H., Zhai, Q., Zhang, J., Wu, X., Sun, J., Xie, Q., Wang, D., and Li, C. (2009). The Arabidopsis RING finger E3 ligase RHA2a is a novel positive regulator of abscisic acid signaling during seed germination and early seedling development. Plant Physiol 150, 463–481.

Burgeff, C., Liljegren, S.J., Tapia-Lopez, R., Yanofsky, M.F., and Alvarez-Buylla, E.R. (2002). MADS-box gene expression in lateral primordia, meristems and differentiated tissues of Arabidopsis thaliana roots. Planta 214, 365–372.

Cai, X.T., Xu, P., Zhao, P.X., Liu, R., Yu, L.H., and Xiang, C.B. (2014). Arabidopsis ERF109 mediates cross-talk between jasmonic acid and auxin biosynthesis during lateral root formation. Nat Commun 5, 5833, doi: 10.1038/ncomms6833.

Cao, M.J., Wang, Z., Zhao, Q., Mao, J.L., Speiser, A., Wirtz, M., Hell, R., Zhu, J.K., and Xiang, C.B. (2014). Sulfate availability affects ABA levels and germination response to ABA and salt stress in Arabidopsis thaliana. Plant J 77, 604–615.

Carles, C., Bies-Etheve, N., Aspart, L., Leon-Kloosterziel, K.M., Koornneef, M., Echeverria, M., and Delseny, M. (2002). Regulation of Arabidopsis thaliana Em genes: role of ABI5. Plant J 30, 373–383.

Chen, H., Zhang, J., Neff, M.M., Hong, S.W., Zhang, H., Deng, X.W., and Xiong, L. (2008). Integration of light and abscisic acid signaling during seed germination and early seedling development. Proc Natl Acad Sci U S A 105, 4495–4500.

Chiang, G.C., Barua, D., Kramer, E.M., Amasino, R.M., and Donohue, K. (2009). Major flowering time gene, flowering locus C, regulates seed germination in Arabidopsis thaliana. Proc Natl Acad Sci U S A 106, 11661–11666.

Dai, M., Xue, Q., McCray, T., Margavage, K., Chen, F., Lee, J.H., Nezames, C.D., Guo, L., Terzaghi, W., Wan, J., Deng, X.W., and Wang, H. (2013). The PP6 phosphatase regulates ABI5 phosphorylation and abscisic acid signaling in Arabidopsis. Plant Cell 25, 517–534.

De Bodt, S., Maere, S., and Van de Peer, Y. (2005). Genome duplication and the origin of angiosperms. Trends Ecol Evol 20, 591–597.

de Folter, S., and Angenent, G.C. (2006). trans meets cis in MADS science. Trends Plant Sci 11, 224–231.

Dekkers, B.J.W., He, H.Z., Hanson, J., Willems, L.A.J., Jamar, D.C.L., Cueff, G., Rajjou, L., Hilhorst, H.W.M., and Bentsink, L. (2016). The Arabidopsis DELAY OF GERMINATION 1 gene affects ABSCISIC ACID INSENSITIVE 5 (ABI5) expression and genetically interacts with ABI3 during Arabidopsis seed development. Plant Journal 85, 451–465.

Du, Z.Y., Chen, M.X., Chen, Q.F., Xiao, S., and Chye, M.L. (2013). Arabidopsis acyl-CoA-binding protein ACBP1 participates in the regulation of seed germination and seedling development. Plant J 74, 294–309.

Feng, C.Z., Chen, Y., Wang, C., Kong, Y.H., Wu, W.H., and Chen, Y.F. (2014). Arabidopsis RAV1 transcription factor, phosphorylated by SnRK2 kinases, regulates the expression of ABI3, ABI4, and ABI5 during seed germination and early seedling development. Plant Journal 80, 654–668.

Finch-Savage, W.E., and Leubner-Metzger, G. (2006). Seed dormancy and the control of germination. New Phytologist 171, 501–523.

Finkelstein, R., Reeves, W., Ariizumi, T., and Steber, C. (2008). Molecular aspects of seed dormancy. Annu Rev Plant Biol 59, 387–415.

Finkelstein, R., Gampala, S.S., Lynch, T.J., Thomas, T.L., and Rock, C.D. (2005). Redundant and distinct functions of the ABA response loci ABA-INSENSITIVE(ABI)5 and ABRE-BINDING FACTOR (ABF)3. Plant Mol Biol 59, 253–267.

Finkelstein, R.R., Gampala, S.S., and Rock, C.D. (2002). Abscisic acid signaling in seeds and seedlings. Plant Cell 14 Suppl, S15-45.

Finkelstein, R.R., Wang, M.L., Lynch, T.J., Rao, S., and Goodman, H.M. (1998). The Arabidopsis abscisic acid response locus ABI4 encodes an APETALA 2 domain protein. Plant Cell 10, 1043–1054.

Fujisawa, M., Nakano, T., and Ito, Y. (2011). Identification of potential target genes for the tomato fruit-ripening regulator RIN by chromatin immunoprecipitation. BMC Plant Biol 11, 26.

Giraudat, J., Hauge, B.M., Valon, C., Smalle, J., Parcy, F., and Goodman, H.M. (1992). Isolation of the Arabidopsis ABI3 gene by positional cloning. Plant Cell 4, 1251–1261.

Gramzow, L., and Theissen, G. (2013). Phylogenomics of MADS-Box Genes in Plants - Two Opposing Life Styles in One Gene Family. Biology (Basel) 2, 1150–1164.

Holdsworth, M.J., Bentsink, L., and Soppe, W.J. (2008). Molecular networks regulating Arabidopsis seed maturation, after-ripening, dormancy and germination. New Phytol 179, 33–54.

Ito, Y., Kitagawa, M., Ihashi, N., Yabe, K., Kimbara, J., Yasuda, J., Ito, H., Inakuma, T., Hiroi, S., and Kasumi, T. (2008). DNA-binding specificity, transcriptional activation potential, and the rin mutation effect for the tomato fruit-ripening regulator RIN. Plant J 55, 212–223.

Jiang, W., and Yu, D. (2009). Arabidopsis WRKY2 transcription factor mediates seed germination and postgermination arrest of development by abscisic acid. BMC Plant Biol 9, 96.

Jiang, Z., Xu, G., Jing, Y., Tang, W., and Lin, R. (2016). Phytochrome B and REVEILLE1/2-mediated signalling controls seed dormancy and germination in Arabidopsis. Nat Commun 7, 12377, doi: 10.1038/ncomms12377.

Kim, J.H., Hyun, W.Y., Nguyen, H.N., Jeong, C.Y., Xiong, L., Hong, S.W., and Lee, H. (2015). AtMyb7, a subgroup 4 R2R3 Myb, negatively regulates ABA-induced inhibition of seed germination by blocking the expression of the bZIP transcription factor ABI5. Plant Cell Environ 38, 559–571.

Kucera, B., Cohn, M.A., and Leubner-Metzger, G. (2005). Plant hormone interactions during seed dormancy release and germination. Seed Sci Res 15, 281–307.

Lee, K., Lee, H.G., Yoon, S., Kim, H.U., and Seo, P.J. (2015). The Arabidopsis MYB96 Transcription Factor Is a Positive Regulator of ABSCISIC ACID-INSENSITIVE4 in the Control of Seed Germination. Plant Physiol 168, 677–689.

Lei, Z.Y., Zhao, P., Cao, M.J., Cui, R., Chen, X., Xiong, L.Z., Zhang, Q.F., Oliver, D.J., and Xiang, C.B. (2007). High-throughput binary vectors for plant gene function analysis. Journal of Integrative Plant Biology 49, 556–567.

Liu, H., and Stone, S.L. (2010). Abscisic acid increases Arabidopsis ABI5 transcription factor levels by promoting KEG E3 ligase self-ubiquitination and proteasomal degradation. Plant Cell 22, 2630–2641.

Liu, X., Hu, P., Huang, M., Tang, Y., Li, Y., Li, L., and Hou, X. (2016). The NF-YC-RGL2 module integrates GA and ABA signalling to regulate seed germination in Arabidopsis. Nat Commun 7, 12768, doi: 10.1038/ncomms12768.

Liu, X.D., Zhang, H., Zhao, Y., Feng, Z.Y., Li, Q., Yang, H.Q., Luan, S., Li, J.M., and He, Z.H. (2013). Auxin controls seed dormancy through stimulation of abscisic acid signaling by inducing ARF-mediated ABI3 activation in Arabidopsis. Proc Natl Acad Sci USA 110, 15485–15490.

Llanes, A., Andrade, A., Masciarelli, O., Alemano, S., and Luna, V. (2016). Drought and salinity alter endogenous hormonal profiles at the seed germination phase. Seed Sci Res 26, 1–13.

Lopez-Molina, L., Mongrand, S., and Chua, N.H. (2001). A postgermination developmental arrest checkpoint is mediated by abscisic acid and requires the ABI5 transcription factor in Arabidopsis. Proc Natl Acad Sci U S A 98, 4782–4787.

Lopez-Molina, L., Mongrand, B., McLachlin, D.T., Chait, B.T., and Chua, N.H. (2002). ABI5 acts downstream of ABI3 to execute an ABA-dependent growth arrest during germination. Plant Journal 32, 317–328.

Meng, L., and Feldman, L. (2010). A rapid TRIzol-based two-step method for DNA-free RNA extraction from Arabidopsis siliques and dry seeds. Biotechnol J 5, 183–186.

Miura, K., Lee, J., Jin, J.B., Yoo, C.Y., Miura, T., and Hasegawa, P.M. (2009). Sumoylation of ABI5 by the Arabidopsis SUMO E3 ligase SIZ1 negatively regulates abscisic acid signaling. Proc Natl Acad Sci U S A 106, 5418–5423.

Nakashima, K., Fujita, Y., Katsura, K., Maruyama, K., Narusaka, Y., Seki, M., Shinozaki, K., and Yamaguchi-Shinozaki, K. (2006). Transcriptional regulation of ABI3- and ABA-responsive genes including RD29B and RD29A in seeds, germinating embryos, and seedlings of Arabidopsis. Plant Mol Biol 60, 51–68.

Nambara, E., and Marion-Poll, A. (2003). ABA action and interactions in seeds. Trends Plant Sci 8, 213–217.

Okamoto, M., Tatematsu, K., Matsui, A., Morosawa, T., Ishida, J., Tanaka, M., Endo, T.A., Mochizuki, Y., Toyoda, T., Kamiya, Y., Shinozaki, K., Nambara, E., and Seki, M. (2010). Genome-wide analysis of endogenous abscisic acid-mediated transcription in dry and imbibed seeds of Arabidopsis using tiling arrays. Plant J 62, 39–51.

Osuna, D., Prieto, P., and Aguilar, M. (2015). Control of Seed Germination and Plant Development by Carbon and Nitrogen Availability. Front Plant Sci 6, 1023.

Parcy, F., Valon, C., Raynal, M., Gaubier-Comella, P., Delseny, M., and Giraudat, J. (1994). Regulation of gene expression programs during Arabidopsis seed development: roles of the ABI3 locus and of endogenous abscisic acid. Plant Cell 6, 1567–1582.

Riechmann, J.L., Wang, M., and Meyerowitz, E.M. (1996). DNA-binding properties of Arabidopsis MADS domain homeotic proteins APETALA1, APETALA3, PISTILLATA and AGAMOUS. Nucleic Acids Res 24, 3134–3141.

Seo, M., Hanada, A., Kuwahara, A., Endo, A., Okamoto, M., Yamauchi, Y., North, H., Marion-Poll, A., Sun, T.P., Koshiba, T., Kamiya, Y., Yamaguchi, S., and Nambara, E. (2006). Regulation of hormone metabolism in Arabidopsis seeds: phytochrome regulation of abscisic acid metabolism and abscisic acid regulation of gibberellin metabolism. Plant J 48, 354–366.

Shu, K., Liu, X.D., Xie, Q., and He, Z.H. (2016). Two Faces of One Seed: Hormonal Regulation of Dormancy and Germination. Mol Plant 9, 34–45.

Shu, K., Zhang, H., Wang, S., Chen, M., Wu, Y., Tang, S., Liu, C., Feng, Y., Cao, X., and Xie, Q. (2013). ABI4 regulates primary seed dormancy by regulating the biogenesis of abscisic acid and gibberellins in arabidopsis. PLoS Genet 9, e1003577, doi:10.1371/journal.pgen.1003577.

Smaczniak, C., Immink, R.G., Angenent, G.C., and Kaufmann, K. (2012). Developmental and evolutionary diversity of plant MADS-domain factors: insights from recent studies. Development 139, 3081–3098.

Stone, S.L., Williams, L.A., Farmer, L.M., Vierstra, R.D., and Callis, J. (2006). KEEP ON GOING, a RING E3 ligase essential for Arabidopsis growth and development, is involved in abscisic acid signaling. Plant Cell 18, 3415–3428.

Tran, L.S., Urao, T., Qin, F., Maruyama, K., Kakimoto, T., Shinozaki, K., and Yamaguchi-Shinozaki, K. (2007). Functional analysis of AHK1/ATHK1 and cytokinin receptor histidine kinases in response to abscisic acid, drought, and salt stress in Arabidopsis. Proc Natl Acad Sci U S A 104, 20623–20628.

Wang, Y., Li, L., Ye, T., Zhao, S., Liu, Z., Feng, Y.Q., and Wu, Y. (2011). Cytokinin antagonizes ABA suppression to seed germination of Arabidopsis by downregulating ABI5 expression. Plant J 68, 249–261.

Winter, D., Vinegar, B., Nahal, H., Ammar, R., Wilson, G.V., and Provart, N.J. (2007). An “Electronic Fluorescent Pictograph” Browser for Exploring and Analyzing Large-Scale Biological Data Sets. Plos One 2: e718. doi:10.1371/journal.pone.0000718.

Xu, D., Li, J., Gangappa, S.N., Hettiarachchi, C., Lin, F., Andersson, M.X., Jiang, Y., Deng, X.W., and Holm, M. (2014). Convergence of Light and ABA signaling on the ABI5 promoter. PLoS Genet 10, e1004197, doi:10.1371/journal.pgen.1004197.

Yan, D., Easwaran, V., Chau, V., Okamoto, M., Ierullo, M., Kimura, M., Endo, A., Yano, R., Pasha, A., Gong, Y., Bi, Y.M., Provart, N., Guttman, D., Krapp, A., Rothstein, S.J., and Nambara, E. (2016). NIN-like protein 8 is a master regulator of nitrate-promoted seed germination in Arabidopsis. Nat Commun 7, 13179, doi: 10.1038/ncomms13179.

Yang, J., Zhang, J., Wang, Z., Zhu, Q., and Wang, W. (2001). Hormonal changes in the grains of rice subjected to water stress during grain filling. Plant Physiol 127, 315–323.

Yu, L.H., Miao, Z.Q., Qi, G.F., Wu, J., Cai, X.T., Mao, J.L., and Xiang, C.B. (2014). MADS-box transcription factor AGL21 regulates lateral root development and responds to multiple external and physiological signals. Mol Plant 7, 1653–1669.

Yu, L.H., Chen, X., Wang, Z., Wang, S.M., Wang, Y.P., Zhu, Q.S., Li, S.G., and Xiang, C.B. (2013). Arabidopsis Enhanced Drought Tolerance1/HOMEODOMAIN GLABROUS11 Confers DroughtTolerance in Transgenic Rice without Yield Penalty. Plant Physiology 162, 1378–1391.

Zhang, H., Cui, F., Wu, Y., Lou, L., Liu, L., Tian, M., Ning, Y., Shu, K., Tang, S., and Xie, Q. (2015). The RING finger ubiquitin E3 ligase SDIR1 targets SDIR1-INTERACTING PROTEIN1 for degradation to modulate the salt stress response and ABA signaling in Arabidopsis. Plant Cell 27, 214–227.

Zhang, Y., Yang, C., Li, Y., Zheng, N., Chen, H., Zhao, Q., Gao, T., Guo, H., and Xie, Q. (2007). SDIR1 is a RING finger E3 ligase that positively regulates stress-responsive abscisic acid signaling in Arabidopsis. Plant Cell 19, 1912–1929.

